## supplement for "The response of Dual-leucine zipper kinase (DLK) to nocodazole: evidence for a homeostatic cytoskeletal repair mechanism"

### RSeQC: Infer Experiment

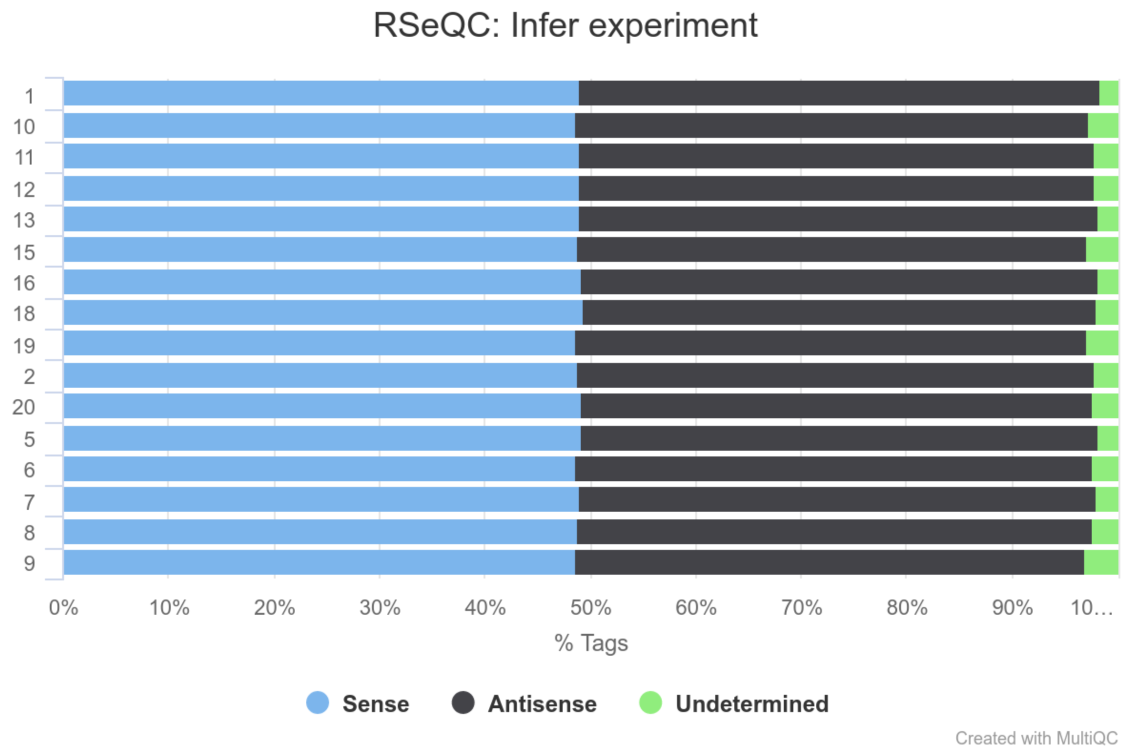


The libraries are unstranded

### FastQC: GC Content (trimmed)

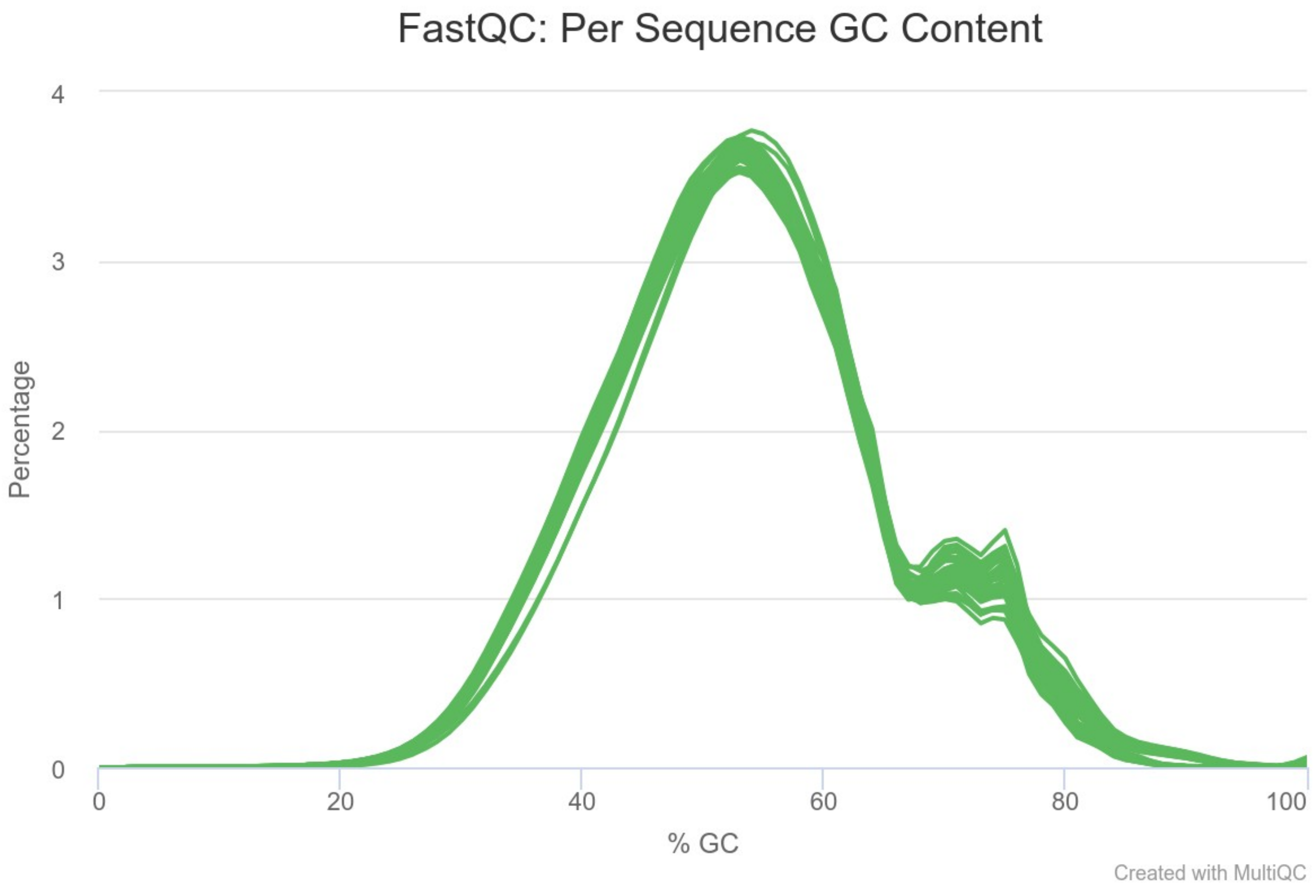

### FastQC: Sequence Quality (trimmed)

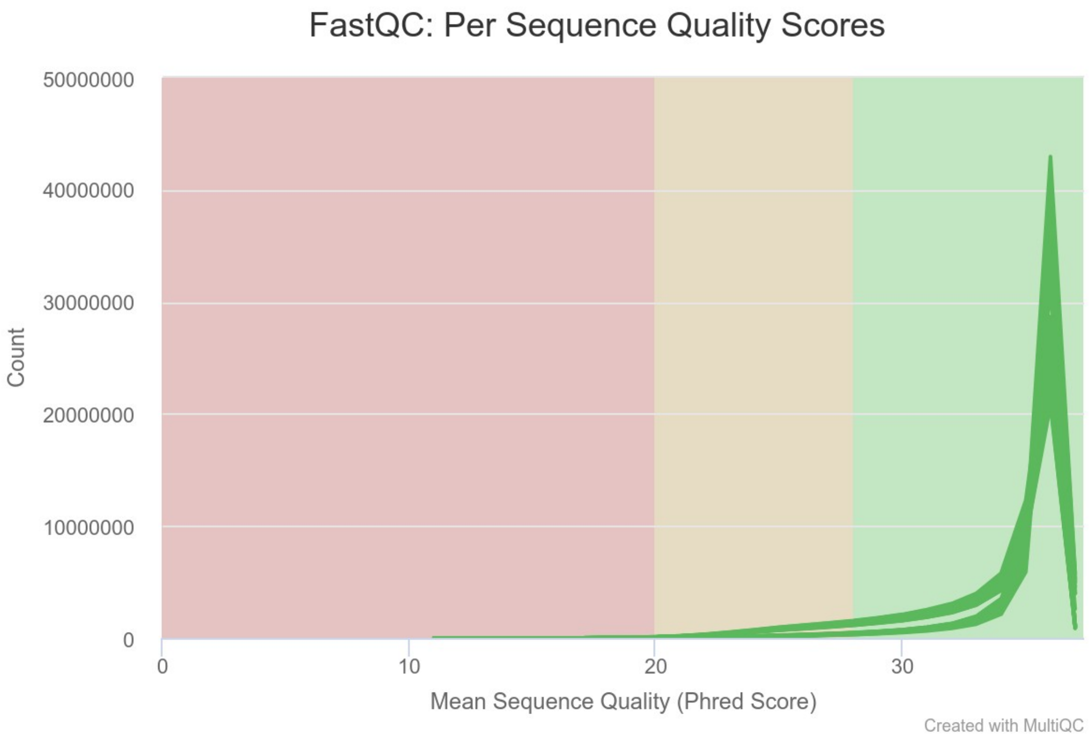

### FastQC: Mean Base Score (trimmed)

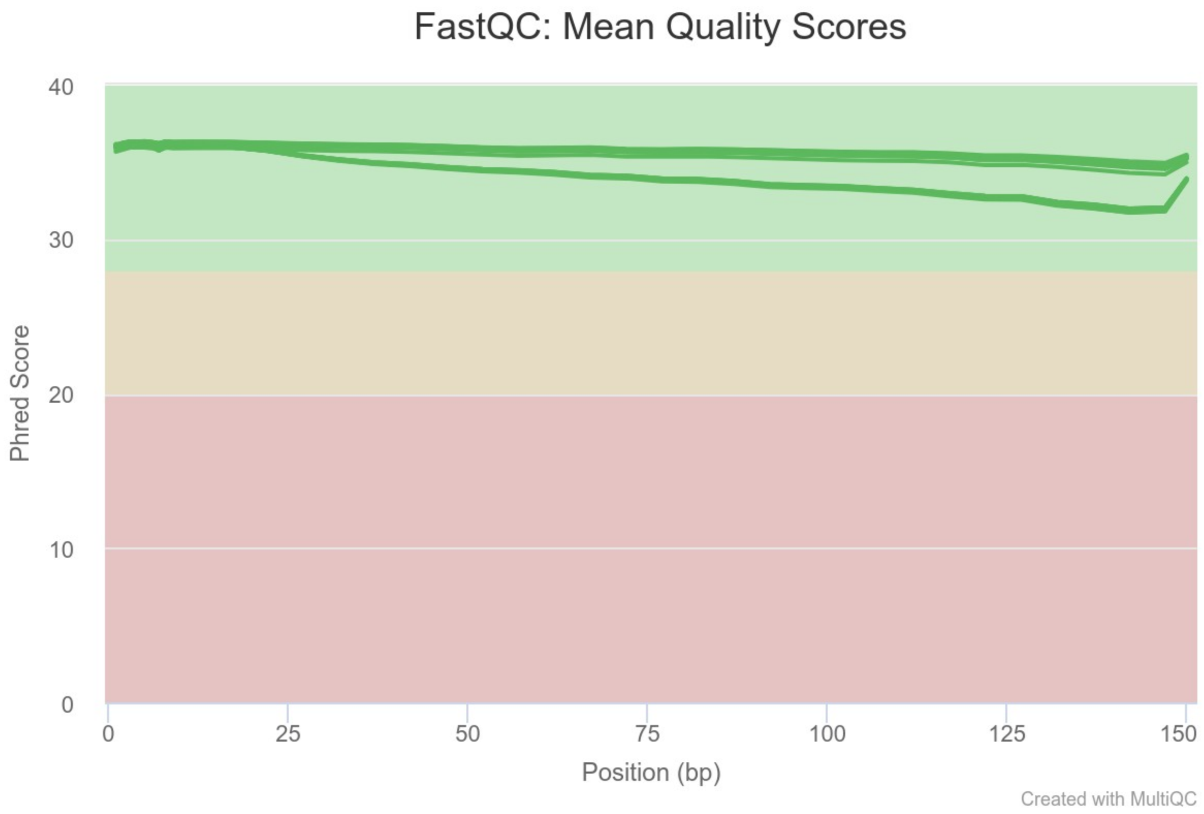

### Picard: Duplication (percent of library)

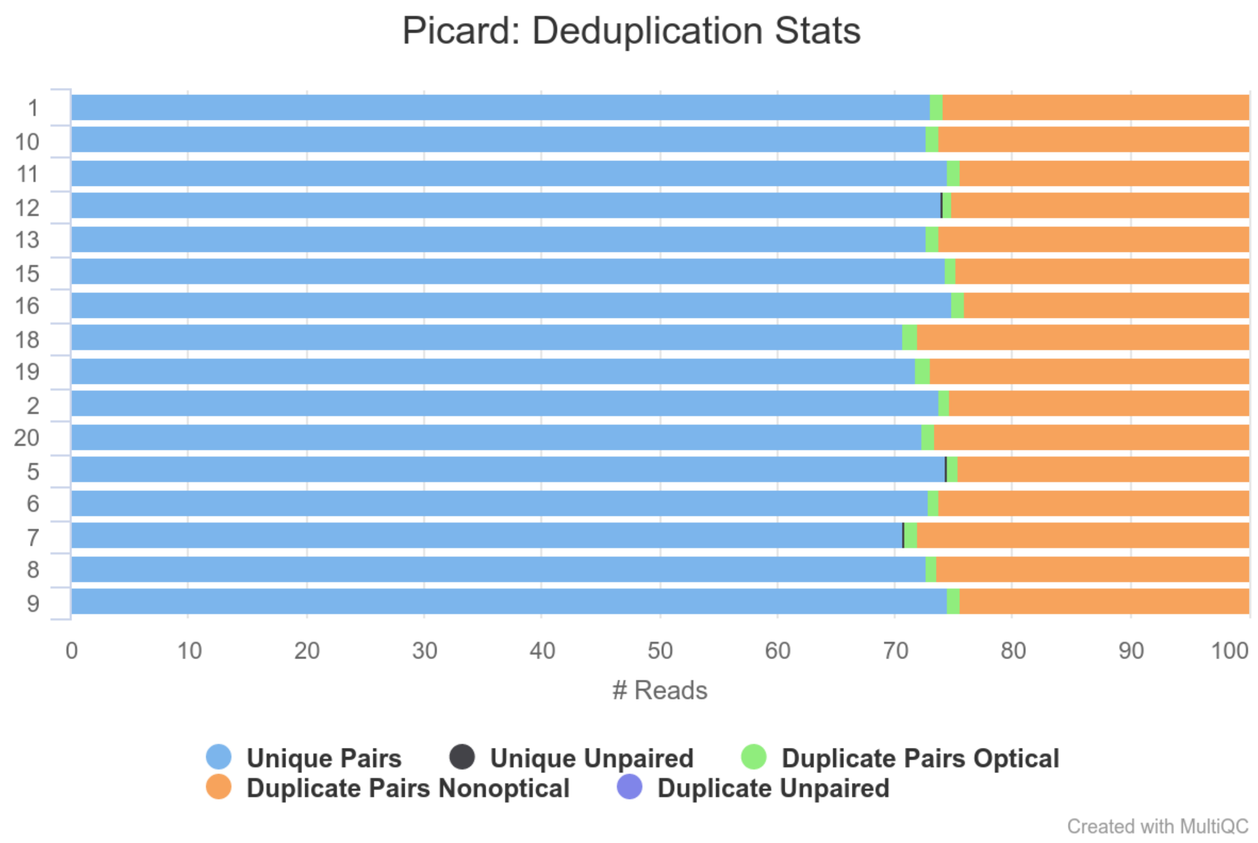

### RseQC: Feature Distribution (percent of library)

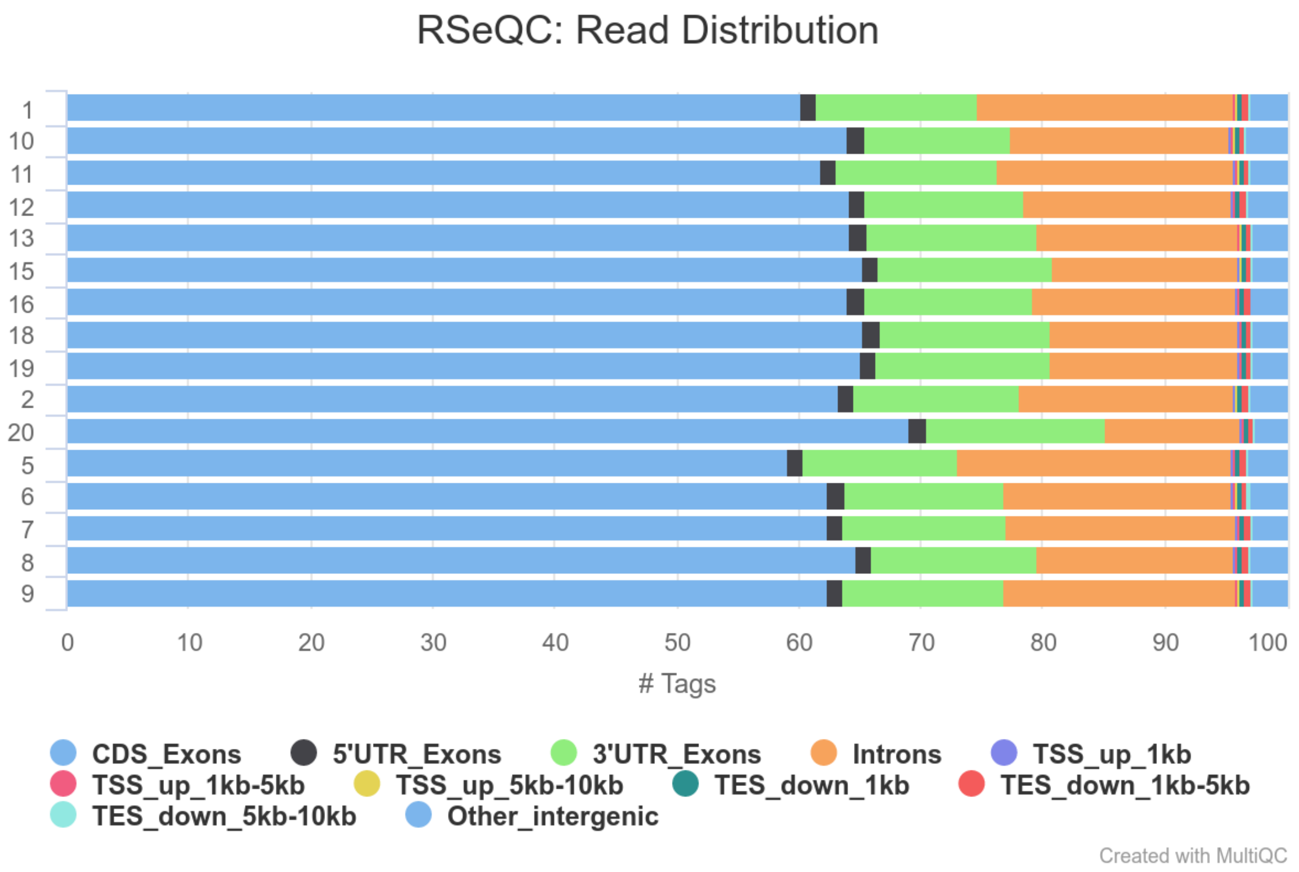

### RseQC: Feature Distribution (raw counts)

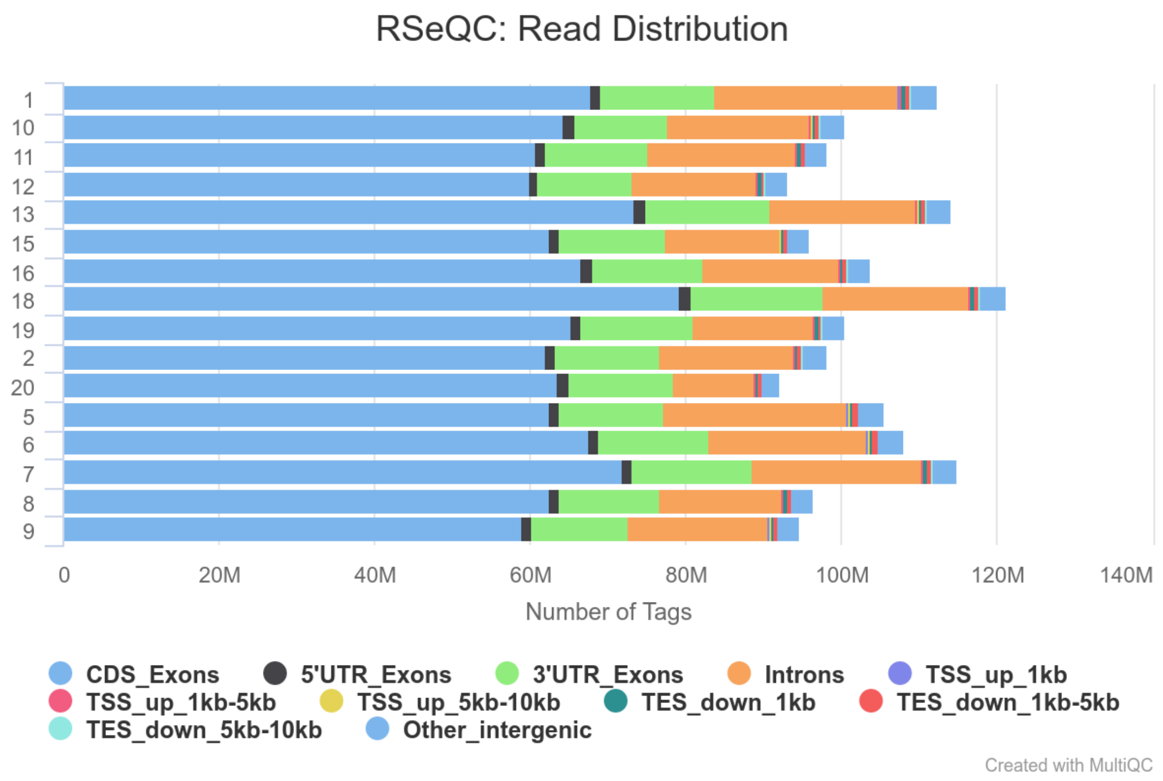

### RseQC: Splice Junction Coverage (percent of library)

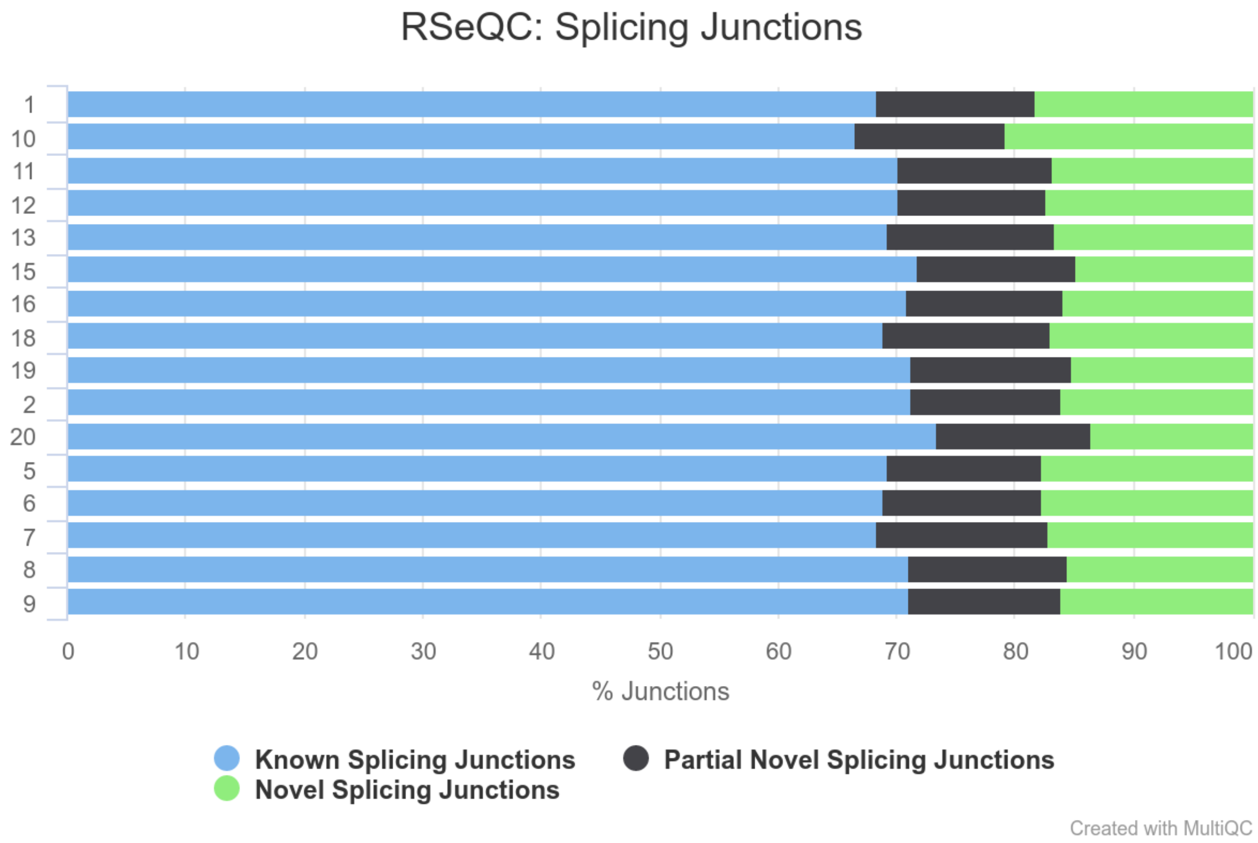

### Qualimap: Gene Body Coverage

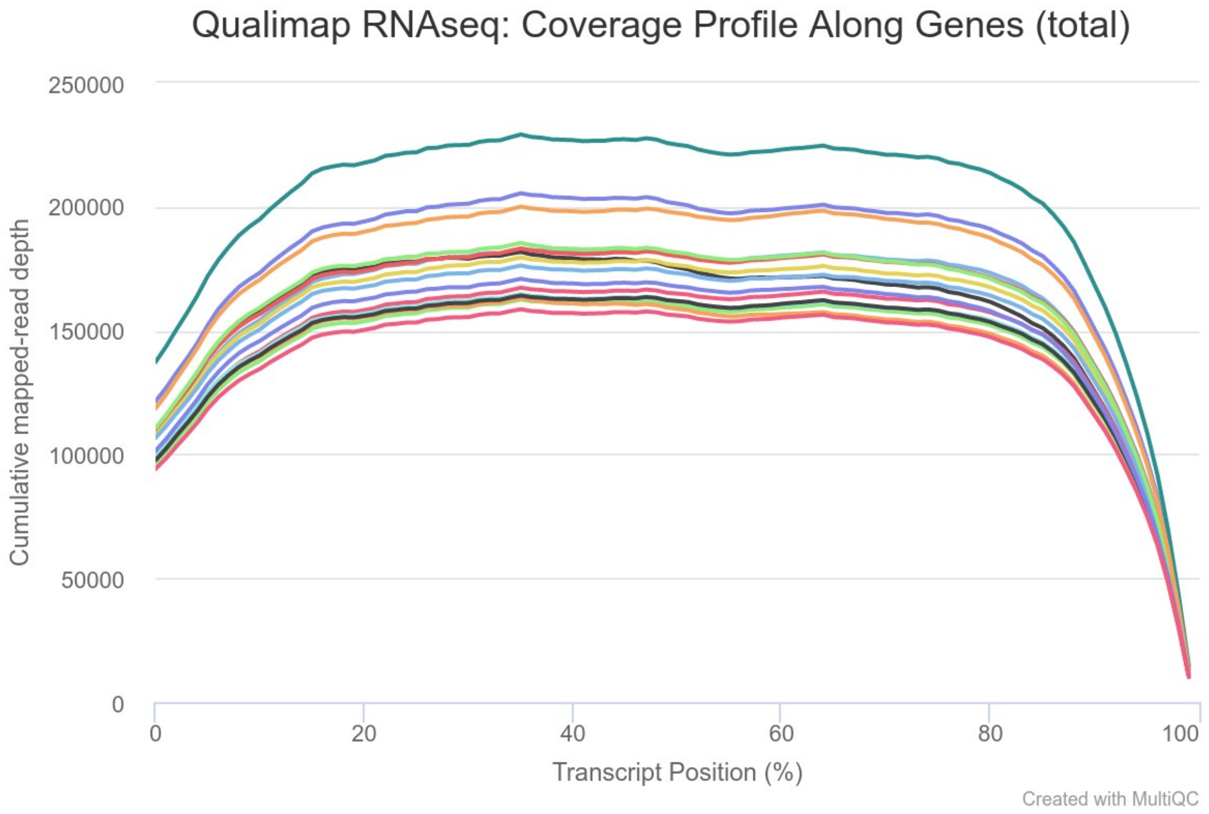
